# Data Independent Acquisition Pipeline for Microbiome Samples (Microbe-DIA)

**DOI:** 10.64898/2026.07.13.738261

**Authors:** Samantha Obermiller, Mary Lipton, Paul Piehowski, Victoria Prozapas, Aivett Bilbao, Lee Ann McCue, Geremy Clair, Isaac Kwame Attah

**Affiliations:** Environmental Molecular Sciences Laboratory, Pacific Northwest National Laboratory, Richland, WA 99354, United States; Biological Sciences Division, Pacific Northwest National Laboratory, Richland, WA 99354, United States

**Keywords:** Data-independent analysis, DIA, microbial communities, metaproteomics, DIA-NN, microbiome

## Abstract

The functional complexity inherent in microbiomes complicates analytical approaches aimed at defining phenotype. As proteins are the functional effectors of microbiome phenotypes, improving the performance of mass spectrometry-based metaproteomics is critical to achieving the functional characterization of these systems. Data-independent acquisition (DIA) improves protein coverage and reduces data missingness when compared to data-dependent acquisition (DDA) in metaproteomics. However, the application of DIA to complex microbial systems remains constrained by analytical throughput and computational scalability. Here, we optimized LC–MS/MS acquisition parameters for both DDA and DIA using a model microbiome, demonstrating how DIA enables increased sample throughput without compromising quantitative performance. In addition, we demonstrated a computationally efficient, library-free DIA workflow that overcomes reliance on empirical spectral libraries. Our analytical and computational innovations establish a scalable and cost-effective pipeline for metaproteomics of complex microbial communities.

## Introduction

Environmental microbiome samples possess a highly complex proteomic profile. This complexity arises from two sources: variation in the abundance of the microbial species as well as variation in the abundance of the proteins in each of the microbes. Further, each microbial genome typically contains thousands of protein-coding genes which are highly homologous across microbes, creating redundantly expressed proteins and peptides. Metaproteomics aims to detect and quantify the suite of proteins expressed in microbial communities, contributing to our understanding of critical functional modulations in these systems and enabling comprehensive multi-omics integration [1].

Liquid chromatography tandem mass spectrometry (LC-MS/MS) proteomic analysis focuses on the evaluation of tryptic peptides from samples where fragmentation of intact peptide precursors is necessary for peptide sequence identification. The traditional approach to fragmentation is data-dependent acquisition (DDA), where narrow mass-to-charge (m/z) windows are used to sequentially isolate the most abundant peptide precursors for MS/MS analysis. This strategy simplifies the analysis of the collected fragment spectra and increases the confidence in peptide identifications. However, because of the limited duty cycle of mass spectrometers, this strategy can acquire spectra for only a limited number of precursor ions, limiting peptide identifications and typically yielding low protein sequence coverage in complex samples, such as microbiomes. Additionally, accurate quantification requires the measurement of peptide intensities at multiple retention times to determine the area under the extracted ion chromatogram (XIC). Since DDA typically avoids fragmenting the same parent ion multiple times, this precludes the use of fragmentation spectra for quantification. In addition, traditional DDA struggles with high data missingness due to this stochastic sampling strategy [2]. Decreasing missing values in LC-MS/MS can provide enlightening functional information for microbiome samples by increasing the number of quantified proteins, especially for the low-abundance precursors [3].

In contrast to DDA, data-independent acquisition (DIA) fragments all precursors within wider m/z windows, enabling more comprehensive sampling of the peptide ion population [4]. In DIA, these windows are fragmented sequentially and cyclically to cover the desired mass range. By forgoing peak selection, DIA yields more reproducible data that includes fragments even from low-abundance precursors [5]. The m/z windows used for fragmentation can vary in width, with researchers weighing the complexity of overlapping peptide fragmentations against the increase in potential peptide identifications. The fragments themselves can also be used for quantification since the same precursors are fragmented multiple times, potentially improving quantification accuracy [6,7,8].

While data-independent acquisition reduces missing values and improves protein quantification, the resulting spectral complexity requires an additional layer of information for processing via a DDA-generated library or a library-free approach [2]. Generating both DDA and DIA data is resource-intensive, and library-free approaches are typically limited in feasibility due to the size of the search database prohibiting the use of large metaproteomes [9]. Previous studies have already compared various DIA software (such as DIA-NN, MaxDIA, and Spectronaut) and found that DIA-NN’s peak selection algorithm is highly effective [10,11,12,13,14]. A recent study by Rajczewski *et al.* also evaluated the reproducibility of DDA and library-free DIA across multiple laboratories using modeled microbiome samples [9]. While they confirmed DIA’s reproducibility and accuracy, their focus on inter-laboratory performance precluded systematic optimization of acquisition parameters. They also noted that further investigation was needed into the scalability of library-free processing to large metaproteomes.

Here, we address these challenges by developing a scalable, cost-effective pipeline for metaproteomic analysis of microbiomes (Figure 1). Building on previous studies, we systematically optimized LC-MS/MS parameters for both DDA and DIA and evaluated multiple library processing strategies using a controlled model microbiome system. We demonstrated that DIA-NN’s library-free mode [12] is currently the most efficient method of analysis while still accurately characterizing the model system. Critically, we present <u>M</u>etaproteomic <u>A</u>nalysis with <u>D</u>IA Library-free (MADL), a two-stage approach that enables library-free DIA processing against a ∼500,000 protein-coding metaproteome database, overcoming the primary computation limitation of this approach.

**Figure 1.**
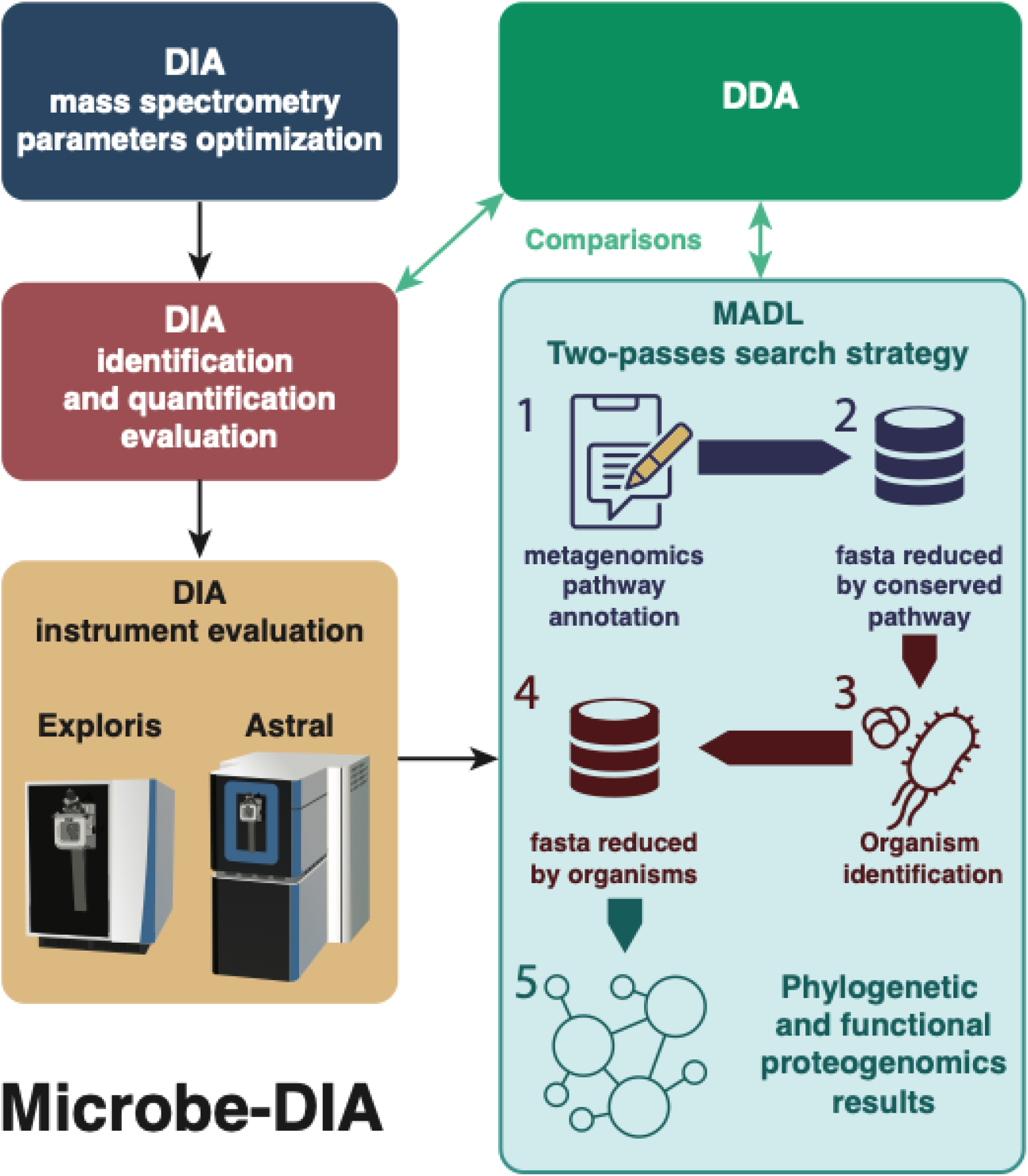
Graphical abstract of the experimentation described in this paper, including the optimization of LC-MS/MS parameters, instrumentation, and analysis. These results were used to develop and validate the Metaproteomic Analysis with DIA-NN Library free (MADL) pipeline.

## Methods

### Designing the model sample and metaproteome

Each of the potential 46 strains of bacteria were cultured individually and underwent genome sequencing [15]. The resulting FASTA files were then used to assess the relatedness of the available species and ensure the model communities simulate microbiome diversity. The proteins for each strain underwent a simulated trypsin digestion, cleaving after arginine or lysine excluding instances followed by proline, allowing for up to 2 missed cleavages and a minimum string length of 10. The resulting fragments were combined for all strains within a species and deduplicated. Relatedness of a species was then calculated as the percent of its peptide fragments shared by any of the other species (Supplementary Figure 1). Five, 24-strain mixtures were designed with species of varying relatedness and informed our comparison of MS1 and MS2 window sizes for DIA and fractionation for DDA libraries (Figure 2, Supplementary Figure 2, Supplementary Table 1). For Figure 1, only Mix1 of the 24-strain dataset was utilized. This decision was based on the consistency across mixtures observed in Supplementary Figure 2. By focusing exclusively on Mix1, the figure provides informative comparisons including MS window size. Two 46-strain mixtures, Mix1 and Mix2, were designed using all available species and informed all further analyses (Figures 3-6).

**Figure 2.**
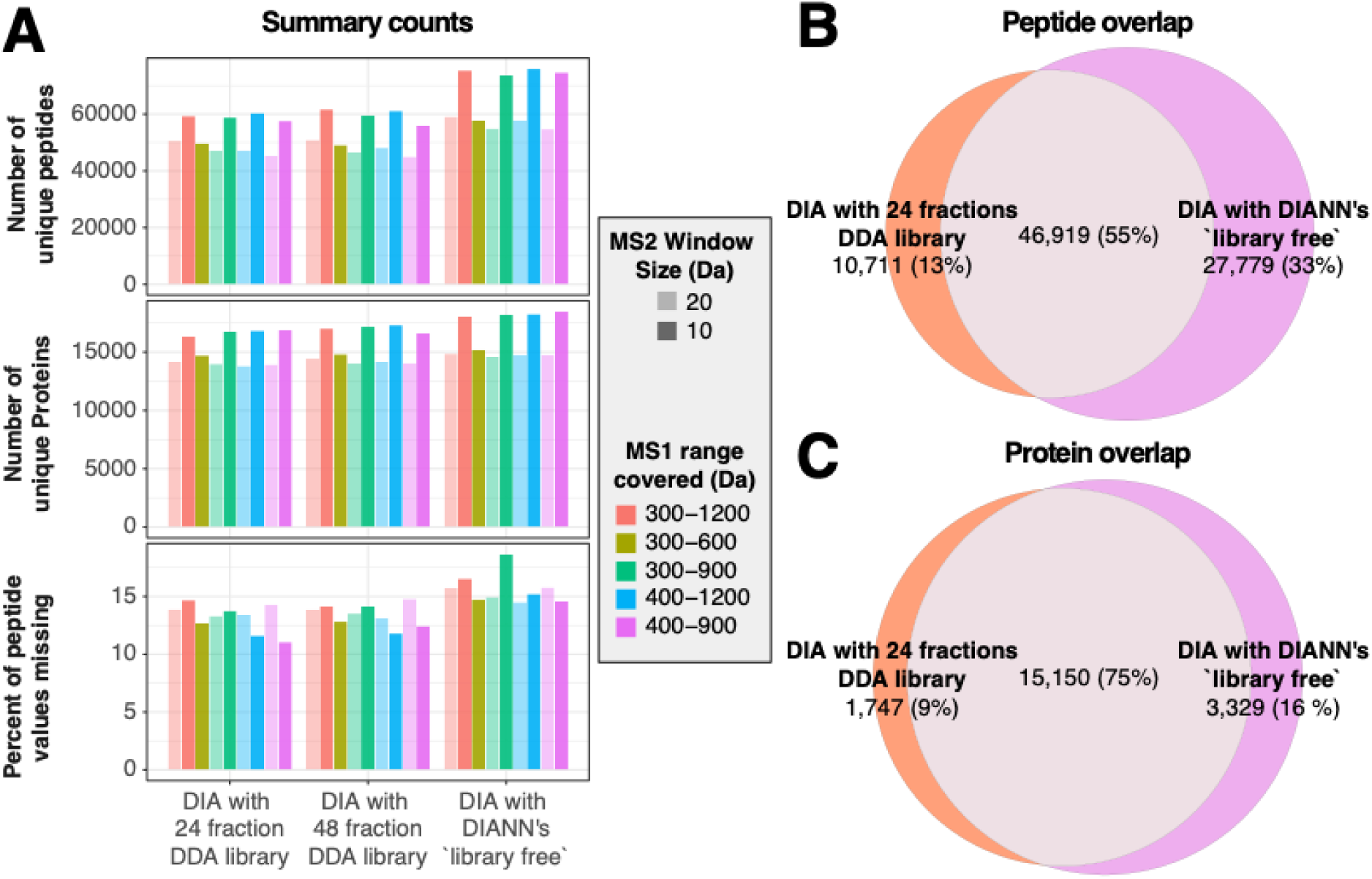
Comparison of DIA window schemes. **(A)** Mix1 of the 24-strain dataset was analyzed by DIA using different MS1 m/z ranges (indicated by bar color), MS2 window sizes (transparent and solid bars), and library types. Results indicate the number of peptide and protein identifications mostly depends on the MS2 window size and using a DDA library versus the in-silico library-free method. Smaller MS2 window sizes and library-free searches yield more detections. Venn diagrams show the overlap in peptide **(B)** and protein identifications **(C)** when DIA data is processed with a DDA library (orange) versus a library-free approach (purple).

**Figure 3.**
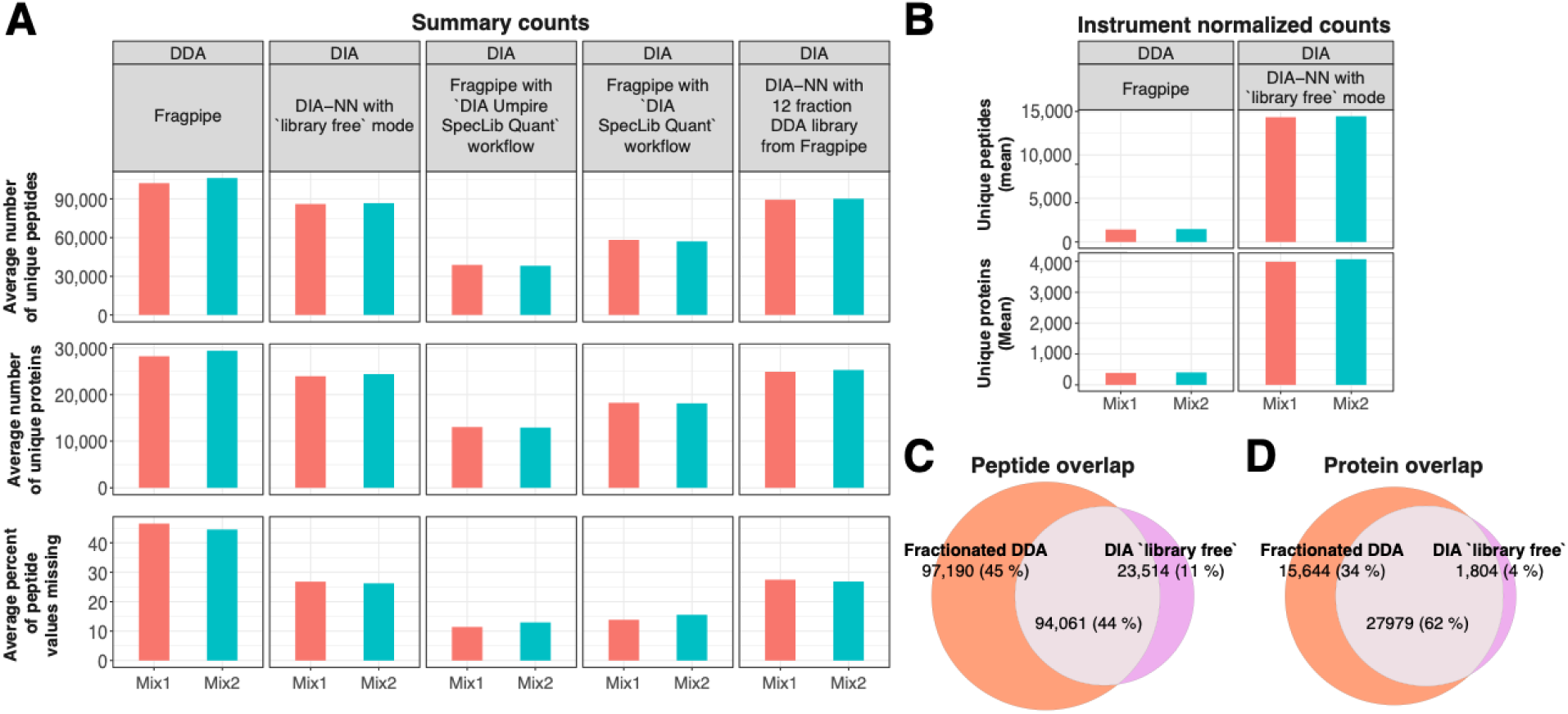
Comparison of DDA and DIA identifications in Mix1 (red) and Mix2 (blue) of the 46-strain dataset using various software tools. **(A)** The number of peptide and protein identifications is lower for DIA than DDA, but DIA has lower percentages of missing peptide values. Amongst the DIA methods, processing DIA with the 12 fraction DDA library provides nearly the same number of identifications as library-free. **(B)** Comparing DDA to the best-performing DIA method, library-free has nearly double the efficiency of DDA when normalized for instrument run time (6 hours per mix vs. 72 hours per mix). **(C)** There is also general agreement amongst the Mix1 peptide and protein identifications made by DDA (orange) and the DIA library-free approach (purple).

### Sample preparation for proteomics analysis

The washed cell pellets were lysed on a bead mill homogenizer (Bead Ruptor Elite) under settings: Speed 6.3 for 45 seconds. Lysates were then centrifuged at 3220 x rpm in large centrifuge for 10min at 4C to remove debris. The supernatant were then transferred to fresh tubes with urea to achieve a final concentration of 8M. A BCA assay (Thermo Fisher Scientific) was then used to estimate protein concentration for downstream processing. Proteins were then reduced by incubating at 10 mM dithiothreitol (DTT) at 37 °C for 1 hr. Lysates were then diluted 8-fold to reduce the urea concentration prior to digestion with trypsin at a 1:50 enzyme:substrate ratio, incubating at 37 °C for 3 hrs. The digestion reaction was then quenched by plunge freezing in LN_2_. Peptides were then desalted using strata C18 cartridges (Phenomenex). Eluents were dried in a speed vac and rehydrated in 50 µL of 18 MΩ water. The concentration of peptides was measured by BCA assay and diluted to 0.1 µg/µL for downstream sample creation (Supplementary Table 1 and 2). The resulting peptide samples were then mixed as described in Supplementary Tables 1, 2, and 3 to create the model microbiomes for our studies.

### LC-MS/MS Analysis

All peptide sample analysis were performed using a Vanquish Neo liquid chromatography system (Thermo Scientific). For longer reverse phase peptide analysis (∼ 9 samples per day (SPD)), the Vanquish Neo LC system was set up with a single pump for both sample trapping and reverse-flow elution onto the analytical column, in the Trap and Elute mode. A 4-cm x 100 um i.d. fused silica column packed with 5µm Jupiter C2 in-house was used for sample trapping. For peptide separation, a fused silica column with an integrated emitter (30 cm × 75 μm i.d.) was packed in-house with 1.7 μm particle size of Waters Acquity BEH particles, Waters). Mobile phases used for sample analysis consisted of (A) 0.1% formic acid in water and (B) 0.1% formic acid in acetonitrile. The 9 SPD method, using a flow rate of 0.20μL/min, has an active gradient of 120 minutes and used the gradient profile; (min, %B): 0.0, 1.0; 11.0, 8.0; 86.0, 21.0; 106.0, 28.0; 116.0, 37.0; 121.0, 75.0; 124.0, 95.0; 129.0, 95.0, Column Wash; 130.0, 50.0; 131,0, 1.0; 132.0, 95.0; 133.0, 1.0; 149.0, 1.0.

For shorter reverse phase elution analysis (∼50 SPD), the Vanquish Neo LC set up similarly, used a PepMap Neo Trap Cartridge (Thermo Scientific) for sample trapping, and a PepMap ES906 analytical column (Thermo Scientific). The 50 SPD method has a 23-minute active gradient, with 6 munities for sample loading and column equilibration. Below is the gradient profile used for this method; (min, Flow Rate, %B): 0.0, 3.50,1.0; 0 .001, 2.80, 1.0; 1.001, 1.30, 4.0; 2.000, 0.800, 8.0; 18.001, 0.800, 22.5; 22.001, 0.800, 35.0; 23.001, 1.10, 45.0; Column Wash; 23.501, 2.80, 99.0; 24.701, 99.0; 25.701, 3.50, 1.0.

The 9 SPD LCMS analysis were performed using the Orbitrap Exploris 480 mass spectrometer, with the analytical column interfaced using a nanoflex nano-electrospray ionization source. The ion transfer tube was set at 300 °C, and the electrospray voltage was 2.2 kV. Sample analysis on the Orbitrap Exploris 480 mass spectrometer was performed either in the data-dependent (DDA) or data-independent (DIA) mode, acquiring data for 120 minutes in both cases. For DDA analysis, full scans were acquired at a resolution of 60k in a scan range of 300 – 1800 m/z. The automatic gain control (AGC) target was set to standard, and maximum IIT set to auto. The Data Dependent Mode was set to Number of Scans, analyzing the top 20 peptide ion peaks per full scan. Peptide ions selected for MS2 analysis were fragmented using higher-energy collision dissociation (HCD) at a normalized collision energy of 30%. The isolation window width for isolating precursor ions was set to 0.7 m/z, with isolation offset set to "off". Resultant fragment ions were detected in the orbitrap, at a resolution of 30K, with first mass set to 110 m/z. The normalized AGC target was set at 250%, and the maximum injection time mode was set to “Auto”. The instrument was set to perform dependent scan on a single charge state precursor, after which the selected peptide is excluded for 45 s. For DIA analysis, the full scan spectra was acquired at a resolution of 120k, over the mass range of 300 – 900 m/z, using a normalized AGC target value of 300% and the and maximum IIT set to auto. DIA mode MS^2^ was performed over the same mass range, using an isolation window size of 10 m/z, window overlap set to 0, and window placement optimization enabled. The cycle time was set to 3 seconds. Resultant fragment data was recorded at an orbitrap resolution of 30 k, over the mass range of 200 – 2000 m/z. The normalized AGC target was set at 800%, maximum IIT set to “Auto”. Similarly, HCD was used for fragmentation at a normalized collision energy of 30%.

All 50 SPD LCMS analysis were performed on the Orbitrap Astral mass spectrometer, with the ES906 analytical column interfaced to the mass spectrometer using an EASY-Spray ion source, and data acquired for 22 minutes. The ion source conditions were 2.2 kV and 300 °C for the spray voltage and Ion transfer tube temperature respectively.

The full scan spectra acquired using the Orbitrap analyzer was performed over a scan range of 380 to 980 m/z at the resolution of 240 k. The normalized AGC target was set at 500%, with the maximum injection time set to 5 ms. MS^2^ analysis was performed using the Astral analyzer, with the DIA window type was set to “Auto”, window placement optimization at “On”, window overlap at 0, and isolation window at 2 m/z. The normalized AGC target was set at 500%, with the maximum injection time set to 3 ms. Fragment analysis was performed using HCD at a normalized collision energy of 25%, and the Astral analyzer scan range set to 150 - 2000 m/z. Loop control mode was set to time, with a value of 0.6 seconds.

### Peptide identification for DDA and DIA data

The raw mass spectrometry files were converted into mzML format using Msconvert with ‘peakPicking’ enabled [16]. The DDA runs were processed in Fragpipe (dx.doi.org/10.17504/protocols.io.bp2l6j6ddvqe/v1), with oxidation on methionine and n-terminal acetylation set as modifications and otherwise default settings [17,18,19,20,21,22,23,24,26,26]. The same process is true for DDA library creation. The DIA runs were processed in either Fragpipe or DIA-NN with the same modifications as DDA [12]. In Fragpipe, library-free DIA was performed using DIA_DIA-Umpire_SpecLib_Quant (dx.doi.org/10.17504/protocols.io.x54v9q46ql3e/v1) and DIA_SpecLib_Quant workflows (dx.doi.org/10.17504/protocols.io.261gey7ydv47/v1), using settings default to that workflow. In DIA-NN, a DDA library generated in Fragpipe (dx.doi.org/10.17504/protocols.io.36wgqxkwklk5/v1 & dx.doi.org/10.17504/protocols.io.j8nlkzpjxl5r/v1) was used as well as the library-free workflow (dx.doi.org/10.17504/protocols.io.yxmvm8jr5g3p/v1 & dx.doi.org/10.17504/protocols.io.5qpvoe4mdl4o/v1). In both cases, protein names were set as the inference method and MBR was enabled, otherwise default settings. The first dataset comparing mass spectrometry parameters utilized Fragpipe v22.0 and DIA-NN v1.9.1. The later datasets utilized Fragpipe v23.0 and DIA-NN v2.0.

### Post-processing for search engine outputs

The combined_ion.tsv output for DDA was already filtered to 1% false discovery (FDR) by Fragpipe. The report.parquet output for DIA was filtered to 1% FDR using columns ‘Lib.PG.Q.Value’, ‘Lib.Q.Value’ and ‘Q.Value’ in R, along with the rest of the post-processing [27,28,29,30,31,32,33,34,35,36]. Peptides were then either 1) filtered to proteotypic peptides (aside from duplicate protein sequences within a species, which were collapsed) or 2) assigned according to parsimony assignment logic. Parsimony assignment logic involves assigning peptides to the protein with the highest number of mapped peptides, continuing in this loop until all peptides are assigned [37,38,39]. Peptides mapping to potential contaminants were removed. The charge states and modifications were then collapsed to the maximum value for each peptide, in accordance with DIA-NN’s documentation. The peptide, mapped protein, species and strain information was then used to establish pmartR components [40]. Using pmartR’s workflow, peptide intensities were log2 transformed and median normalized, then protein rollup was performed using the median peptide value for each protein. For the 46-strain dataset, the resulting pmartR objects were then filtered to proteins with a minimum of two non-missing values amongst the three replicates for each of the two mixtures. pmartR’s imd_anova function was then used to perform a two-sample t-test, assuming equal variance between the two mixtures and using a Benjamin-Hochberg FDR adjustment.

### Scaling DIA-NN library-free searches to metaproteomes

We obtained KEGG annotations from BLAST Koala for the proteins found in one of the 46-strain mixtures processed in DIA-NN’s library-free mode with only proteotypic peptides [41,42]. The pathways for those KO annotations were then obtained via the KEGG API [43]. Pathway enrichment analysis using a “holm” adjusted Fisher’s exact test was then performed comparing the pathways associated with the 60 most abundant, strain unique proteins to those associated with all the proteins found in that dataset (Supplementary Figure 3). The 20 most significant pathways were then used to filter the original database as well as three external FASTAs. The external FASTAs were chosen because they had varying sizes and represented varying microbiome communities (gut metaproteome originally sourced from the Human Microbiome Project in 2020 (1,162,845 proteins), a collection of 35 soil isolates generated by the Joint Genome Institute (171,005 proteins) and a soil metaproteome also generated by JGI (2,360,231 proteins) [44,45,46]. Each of these three external databases were then combined with the original and used to re-search the 46-strain DIA dataset in DIA-NN library-free mode. Organism attribution depended on what was provided by the original FASTAs.

### Metaproteomic Analysis with DIA-NN Library Free (MADL)

By generalizing the conserved pathways used for FASTA filtering and enforcing stricter confidence thresholds for organism identification, we built on the testing described in the previous section and developed a refined pipeline for using DIA-NN’s library-free search in metaproteomic samples, described below and in Figure 1. (1) KEGG annotations of the FASTA are performed locally using a fork of the kofam_scan repository available on Github [47]. Associated pathways are obtained via a static Json provided by KEGG [43,48,49]. Organism attributions are again dependent on what’s provided in the FASTA. (2) The annotated FASTA is filtered to a generalized list of pathways determined by previous studies to be conserved across bacterial species [50,51]. After the pathway down-selected FASTA has been run through DIA-NN’s library-free mode, the results are filtered to only proteotypic peptides with an initial FDR threshold of 0.01. A stricter ‘PG.Q. Value’ (protein group false discovery rate) threshold is set by generating a histogram of PG.Q.Values for these peptides (100 bins) and finding the first bin with a large drop in frequency (<0.01% of the largest bar). The new threshold is the max value of the last high-count bin [52–54]. (3) After filtering to peptides with PG.Q.Values less than or equal to new threshold, the number of proteins associated with each organism are counted and only organisms with at least 5 filter passing proteins are considered “detected”. (4) The original FASTA is re-filtered using the list of detected organisms and (5) run through DIA-NN “library free” mode again to obtain phylogenetic and function results. The full pipeline is available at TBD.

### Statistics and Reproducibility

For the primary model microbiome experiments, peptide samples were prepared from mixtures described in Supplementary Tables 1, 2, and 3. Biological replicates were defined as independently processed aliquots of each model microbiome mixture. Technical replicates were not included, as sample processing and LC-MS/MS acquisition were performed once per biological replicate. The 24-strain dataset involved five mixtures with no biological replicates and no downstream statistics beyond peptide and protein counts. The 46-strain dataset had two mixtures, Mix1 and Mix2, with 3 biological replicates each. Statistical analyses were detailed in the relevant Results and Methods subsections. Relevant versioning for R (version 4.4.2) packages includes tidyverse (version 2.0.0), here (version 1.0.1), pmartR (version 2.4.6), microseq (version 2.1.6), arrow (version 18.1.0), data.table (version 1.16.2), readxl (version 1.4.3), readr (version 2.1.5), scales (version 1.3.0), cowplot (version 1.1.3). Relevant versioning for Python (version 3.11.0) packages includes pandas (version 3.0.1), pyarrow (version 23.0.1), ruby (version 3.4.8), hmmer (version 3.4), parallel (version 20170422) and matplotlib (version 3.10.8). Figures in this paper were minimally adjusted using Adobe Illustrator 2026.

## Results

### Simulating microbiome complexity

The goal of this study was to develop and validate a robust workflow using DIA-based proteomics for the characterization of microbiomes. We used mixtures of peptides isolated from axenic cultured microbial species to model microbiome-like complexity, where microbial species composition and concentrations were predetermined. To design these mixtures and to ensure they reflected realistic complexity, we assessed the phylogenetic relatedness of the available species. In total, we used 46 strain isolates from 28 bacterial species originating from a soil microbiome [15]. The full list of strains and their genome characteristics is provided in Supplementary Tables 1 & 2. We utilized previously cultured and sequenced organisms and performed in silico trypsin digestion on their individual FASTA files. In-silico peptide sequences from all strains of each species were merged into single-species level lists. The relatedness of the species was measured as the percent of tryptic peptide sequences shared with any of the other available species (Supplementary Figure 1). This analysis indicated that there is a wide range of relatedness in our pool of organisms. For instance, multiple species in our pool belong to the *Bacillus* Genus; while some have a high degree of genetic overlap (i.e., *Bacillus aryabhattai*), others were less genetically related (i.e., *Bacillus pumilus*). This variation highlights the variability of taxonomic assignment and the importance of independently assessing relatedness. Using this information, we designed five, 24-strain mixtures using the same subset of low, medium, and highly related species (23 species in total, Supplementary Table 1). When multiple strains were available for a given species, one was chosen at random. We also designed two 46-strain mixtures using all available strains to compare DDA and DIA results (Supplementary Table 2). We varied the proportion of each strain between the two 46-strain mixtures, providing ground truth for quantification assessments (Supplementary Table 3).

### DIA window schemes

Our first task was to determine the optimal instrument and search settings to be used for the evaluation of the microbiome model samples. To refine the generation of DIA mass spectrometry data, the 24-strain mixtures were analyzed using various MS1 and MS2 window sizes (Supplementary Table 1). The raw files were processed in DIA-NN using 24- and 48-fraction DDA libraries from Fragpipe as well as DIA-NN’s library-free mode, as described in the methods section [12,17,18,19,20,21,22,23,24,25,26]. In total, the 24-fraction DDA library generated 373,490 peptides while the 48 fraction DDA library generated 435,226 peptides. At this stage, we restricted the data analysis to proteotypic peptides, i.e., experimentally observable peptides that each map to only one protein [55], since the process for assigning redundant peptides varies across the field and can have a large impact on results. The 24-fraction DDA library had 259,787 proteotypic peptides and the 48 fraction DDA library had 308,221 proteotypic peptides.

The DIA datasets with a 10 Da MS2 isolation windows, analyzed with DDA libraries, yielded an average of 57,371 peptide identifications, whereas a 20 Da window yielded 47,627 (Figure 2A). The relatively large difference of about 10,000 peptides indicates the smaller MS2 window size enabled more efficient identification, likely because it produced less complex MS2 spectra [56]. Further, there was relatively low variation in the number of peptides identified across MS1 m/z ranges. This is indicated by the small standard deviation of peptide counts within the 10 Da MS2 window size (4,470 peptides) and 20 Da MS2 window size (2,184 peptides). This result indicates that the MS1 m/z range had little effect on the number of peptides detected. The common *m/z* range of 400-900 is where a large portion of detectable peptides fall due to factors such as tryptic peptide length and ionization efficiency [4,56]. Overall, these results indicate that the number of peptide and protein identifications is more dependent on MS2 window size than m/z range.

We observed a similar number of peptides identified between the two DDA libraries (i.e., 24- and 48-fraction libraries) suggesting that higher levels of fractionation are not beneficial for less complex microbial samples, as increasing the number of fractions did not result in a significant increase in the identified peptides and proteins. Next, we performed a “library-free” search using DIA-NN to evaluate the performance of this algorithm on complex metaproteomes. Using a 10 Da MS2 window, we identified an average of 71,494 peptides, which represents about a 25% increase in peptide identifications without relying on DDA data. Between these two strategies, only 13% of peptides and 9% of proteins were not observed using the library-free mode in place of a DDA library (Figures 2B and 2C). The overlap in DIA identifications between library-free mode and the 24-fraction DDA library suggests that both approaches provide comparable information overall, lending confidence to the newer in silico methods.

### Identification and quantitation

Having identified a suitable DIA instrument method for microbiome applications, the next step was to increase complexity by using the 46-strain mixtures to further investigate various processing methods and compare their results to DDA. DIA data were generated using a 300 - 900 Da MS1 m/z range, 10 Da MS2 window size, and a total of 6 runs (2 mixtures x 3 biological replicates). DDA data were generated using high pH reverse phase fractionation, resulting in 72 total runs (2 mixtures x 3 biological replicates x 12 fractions). Since each fraction was analyzed with a 2-hour gradient, the total analysis time for DDA was 144 hours, while the total analysis time for DIA was 12 hours. The 12 fraction DDA data was also used to generate a library, which had 379,015 total peptides and 271,829 proteotypic peptides. Considering the previous success of DIA-NN’s library-free mode, we evaluated two additional processing methods from Fragpipe that do not rely on DDA data: DIA_DIA-Umpire_SpecLib_Quant and DIA_SpecLib_Quant [16,17,18,19,20,21,22,23,24,25].

The overall number of peptides identified does not appear to vary between the two 46-strain mixtures. DDA combined with fractionation yielded more average peptide (104,199) and protein (28,813) identifications than any of the DIA methods tested, including library-free (86,368 peptides and 24,176 proteins) (Figure 3A). To calculate missingness, we defined the total peptide identification space as the set of all peptides identified across mixtures and replicates for a given acquisition mode (DDA vs DIA) and search method (library-based vs. library-free). We then calculated missingness as the percentage of those peptide values missing for each replicate within that acquisition mode and search method. DDA has the highest percentage of missing peptides in Mix1 (46.6%) and Mix2 (44.5%) when averaged across the three replicates, compared with DIA library-free, which has a 26.8% missingness for Mix1 and 26.3% for Mix2. The average number of peptide identifications from DIA in Fragpipe was 45.4% less than the average for DIA-NN. To simplify future computations, we decided to focus further comparisons on DIA results from DIA-NN.

One consideration in comparing DDA to DIA is the instrument time needed to fully analyze each sample. The DDA data was collected using two-dimensional liquid chromatography (2D-LC) reverse phase analysis to reflect the current approach applied in our lab, while DIA was one-dimensional liquid chromatography (1D-LC) reverse phase analysis. The total analysis time per mixture was 72 hours for DDA and 6 hours for DIA. When normalized for analysis time, the peptide identification yield in DIA mode is nearly 900% higher than in DDA mode (Figure 3B). Importantly, the DDA and DIA library-free data showed substantial overlap in peptide and protein identifications (Figure 3C). Despite identifying fewer proteins than our DDA approach, DIA enables rapid microbiome characterization and can dramatically expand metaproteomics throughput, allowing larger experimental designs across space, time, and cohort size.

Up to this point, our comparisons have focused on qualitative assessment of peptide identifications. However, quantitative change in protein abundance can provide insight into the underlying functional phenotypes in microbial samples. To assess the quantitative performance of DIA and DDA, we tested for differential abundance (T-test with Benjamini-Hochberg FDR adjustment) of each protein. Directionality was determined from the log-transformed fold change between Mix1 and Mix2 in the 46-strain dataset. Since we altered the proportions of the peptides from each strain between Mix1 and Mix2, we expected the protein abundances to differ significantly between the two peptide mixtures. Previously, our assessment only included proteotypic peptides, because non-proteotypic peptides can introduce ambiguity. Here, to evaluate protein-level abundance, we compare a proteotypic-only approach with an approach that uses both proteotypic and non-proteotypic peptides assigned using the parsimony attribution approach [37,38,39]. This is a popular method that assigns peptides to the protein with the highest number of mapped peptides, as detailed in the methods. Importantly, all DIA processing strategies resulted in the detection of more significantly modulated proteins than DDA (i.e., over 14,776 proteins in DIA library-free vs. 7,817 in DDA), indicating improved quantification, likely due to fewer missing peptide values (Figure 4A). Most statistically significant proteins also had the correct fold change direction (Figure 4B). By utilizing the parsimony method, as described in the post processing section of the methods, to incorporate non-proteotypic peptides, we detected 6,824 additional proteins correctly quantified in DIA library-free and 3,954 in DDA. This suggests that strategies for peptide assignments, such as the parsimony method, can further improve quantification. We did not comprehensively evaluate the different peptide assignment methods as this was not the focus of our study. Since the proportions of the strains and species in each mixture are known, we tested if we were able to reconstitute the microbial composition by adding together the protein intensities of each species or strain and calculating it as a proportion of the total intensity for Mix1 (Figure 4C & D). Across all the DIA processing strategies and the DDA data, Mix1 had proteins attributed to nearly all the same species (between 26 and 28 out of 28 species) and strains (between 37 and 44 out of 46 strains). The proportions are also consistent across processing and acquisition strategies, even when they deviate from the known proportions. This holds true with the use of the parsimony assignment method. This analysis demonstrates that DIA matches DDA’s ability to accurately characterize highly complex mixtures, capturing not only protein abundance changes but also species composition.

**Figure 4.**
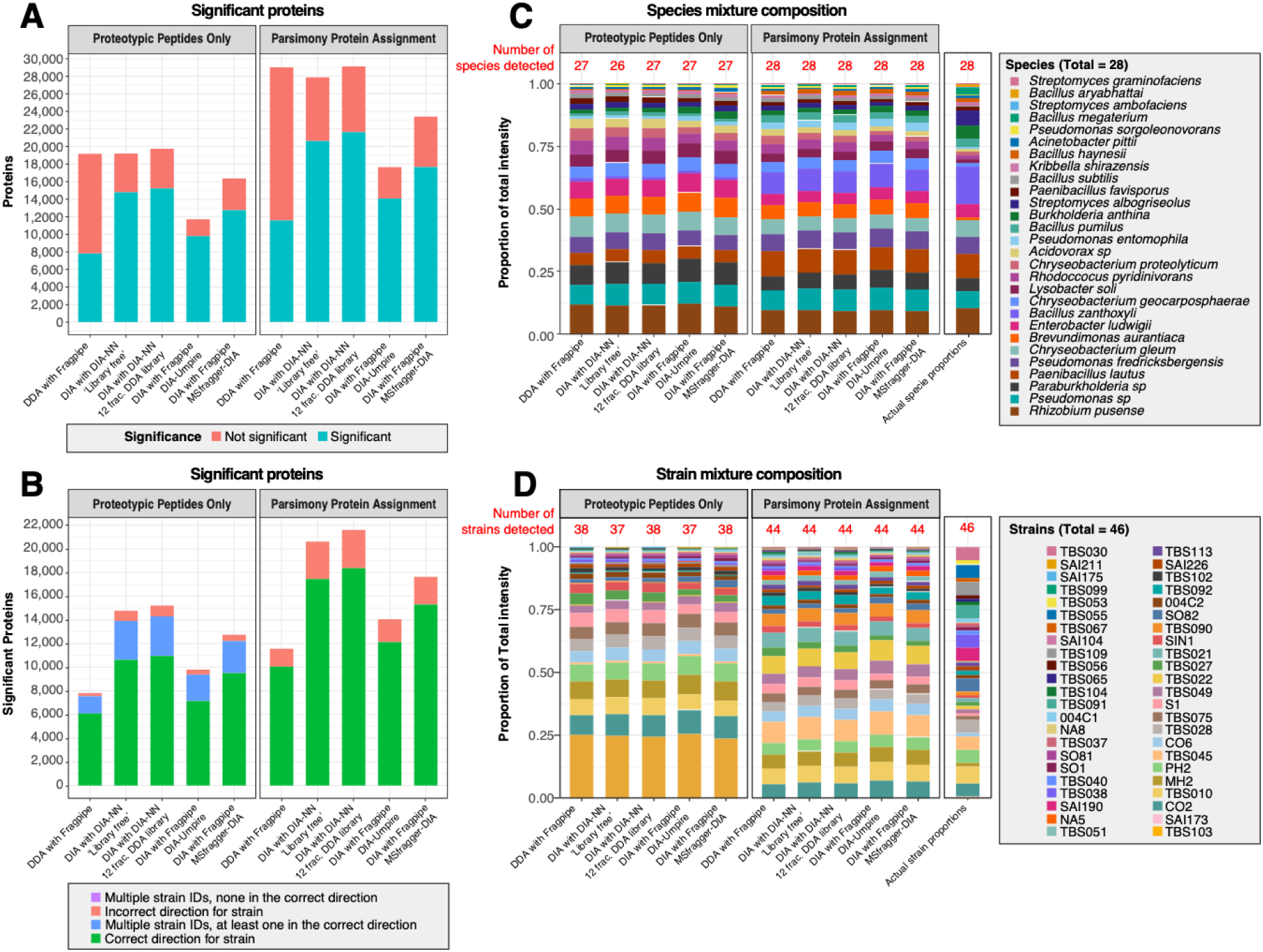
Comparison of DDA and DIA quantification. Quantification of the 46-strain mixtures indicates that **(A)** DIA provides a higher proportion of statistically significant proteins than DDA and **(B)** of those significant proteins, more are correctly quantified in DIA than DDA. Mixture compositions for Mix1 at both the species (**C**) and strain (**D**) level are consistent across DDA and the various DIA processing strategies. These trends hold true both when using only proteotypic peptides for quantification, this study’s default, and when using parsimony protein assignment logic, a common approach in metaproteomics.

The recently released Astral mass spectrometers (Thermo Scientific) can achieve DIA scan rates that exceed 200 Hz with its design incorporating an astral mass analyzer, whereas Orbitrap only instruments are typically limited to 40-60 hertz. Preliminary reports in the literature suggest the Astral will further improve identification efficiency and quantitative accuracy [57]. Therefore, we set out to benchmark this instrument against our model system and the processing approach described above. Each 46-strain mixture was analyzed using DIA mode on the Astral instrument (3 biological replicates, each run had an analysis time of 21.88 minutes) for a total of 1.1 hours. Using DIA-NN’s library-free mode and only proteotypic peptides, we report that the Astral 1-hour analysis time yielded 12,877 more peptide and 1,017 protein identifications, averaged across mixtures, than the Exploris 6-hour analysis time (Figure 5A). The Astral also provides a similar number of correctly quantified proteins (10,506 proteins for the Astral vs. 10,624 for the Exploris), indicating that protein quantification is not impacted by the reduced run time on the Astral (Figure 5C). Normalizing for instrument run time, this is a 468% improvement in hourly protein identification (Figure 5B).

**Figure 5.**
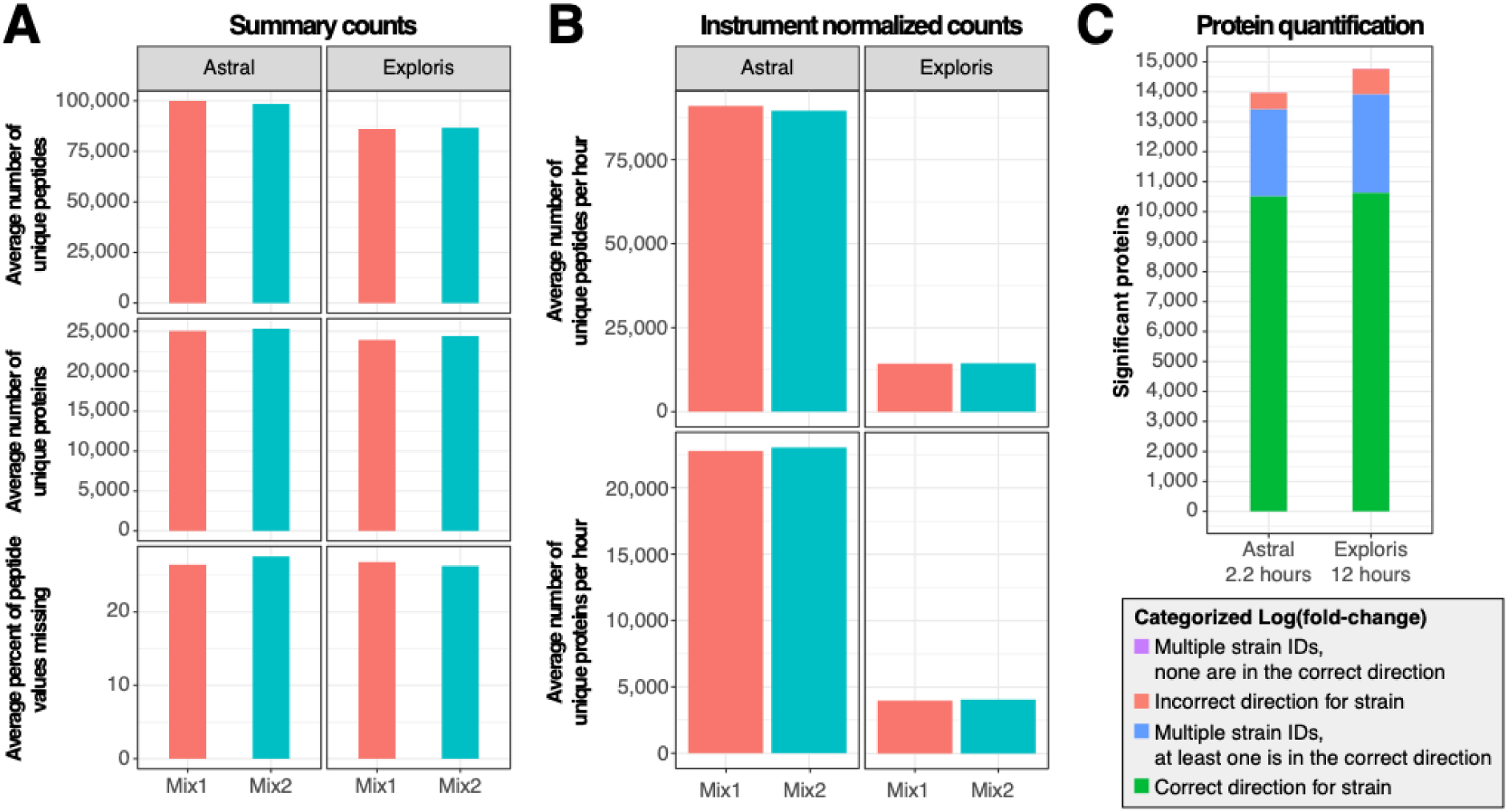
Comparison of the Astral DIA and the Exploris DIA results on Mix1 (red) and Mix2 (blue) from the 46-strain dataset. **(A)** The Astral only detects slightly more peptides and proteins than the Exploris, and **(C)** both instruments correctly quantify a similar number of proteins, but when **(B)** normalized for instrument time (1 hour per mix for the Astral vs. 6 hours per mix for the Exploris), the Astral provides about 9-times more peptide and protein identifications per hour than the Exploris.

### Computational metagenomic approaches

Having confirmed that DIA library-free is a highly effective alternative to a DDA data-generated library, we next need to assess whether it can be used when searching against the large metagenomes typical of natural microbiomes, which range in microbial diversity from 100’s (gut microbiome) to 1000’s (soil microbiomes) of different species. We attempted running library-free on the 46-strain dataset using an external soil metaproteome (2,360,231 proteins) amended with the metaproteome from our 46 strains (248,658 proteins), but DIA-NN failed to complete the search and did not provide an error log file or any other output. We determined that a new approach was necessary for running DIA-NN library free on microbiome samples, both due to this run failure and because searching a metaproteome of that size has been known to produce false discovery issues even if completed [58].

The main goals of metaproteomics are (i) to identify which organisms are present and actively functioning in a community, and (ii) to determine what functions are performed by this community. To fulfill the goals of metaproteomics in a computationally efficient fashion, we developed a two-pass search approach which uses DIA-NN’s library free mode: the first pass is used to identify the organisms potentially present in the samples, and the second pass is used for the functional characterization of the microbiome. We termed this approach Metaproteomic Analysis with DIA-NN Library-free (MADL). Previous work has demonstrated that multiple pass approaches can improve the performance of large database searches [58]. MADL builds on this concept by improving the stepwise logic and imposing strict FDR thresholds for organism identification. The full MADL pipeline was deposited on Github at TBD, allowing people to reproduce our approach.

In the first pass of MADL, we subset the search database to proteins from conserved pathways. Most microorganisms conserve essential processes, such as glycolysis, TCA cycle, ATP generation, ribosomal replication, or DNA replication. While core metabolic and other essential functions are shared across species, evolutionary pressure drives discrepancies in the genes and proteins involved. To test the viability of MADL’s first pass, we needed to confirm that these essential processes exist for the organisms in our dataset and that they will reduce the search databases to a size reasonable for DIA-NN library free. To confirm that core processes are conserved in our 46-strain dataset, we performed KEGG pathway enrichment analysis comparing pathways for the 60 highest intensity proteins per strain, to all pathways identified in the dataset. As anticipated, the enriched pathways comprised many core functions, including “Glycolysis/Gluconeogenesis”, “TCA cycle”, “Microbial metabolism in diverse environments”, “Ribosome”, or “Nucleotide metabolism” (Supplementary Figure 3A). We also confirmed that these enriched pathways were specific enough to reduce both our FASTA and three external metaproteomes of varying size, as detailed in the methods (Supplementary Figure 3B). Note that a large portion of the decrease in size came from whether we were able to attribute a Kegg Ontology (KO) to the protein sequences present in the FASTA.

Having confirmed the viability of our first pass strategy, we next needed to remove bias towards our test dataset and validate the full MADL pipeline. We therefore performed the full two-pass search, as depicted in Figure 1, using our FASTA and the three external metaproteomes. (1) We annotated all three test FASTA databases (three external metaproteomes, each with our original database appended) and (2) filtered to proteins sequences associated with pathways previously reported as broadly conserved across bacterial species, as described in the methods section [50,51]. (3) Each of these pathways filtered databases were then used to reprocess the 46-strain dataset in DIA-NN library free mode. To generate a list of ‘detected’ organisms for each set of results, identifications were first restricted to proteotypic peptides with an initial FDR of 0.01. Histograms of DIA-NN protein group confidence values (PG.Q.Value) were then generated after applying a - log10 transformation. To remove low confidence tails, each set of results received a new PG.Q.Value threshold defined as the upper boundary of the last histogram bin whose frequency was at least 0.01% of the maximum bin height. (4) Each test FASTA was then filtered to its list of detected organisms and (5) used to re-process the 46-strain dataset in DIA-NN, obtaining taxonomic and functional information on the sample.

The generalized set of pathways successfully reduced the test FASTAs to a size tractable for DIA-NN library-free processing (e.g. about 250,000 sequences) and successfully enabled detection of 25 of the 28 known organisms across all three test FASTAs (Figure 6 A, B & C). Based on the results of the two-pass pipeline, the number of falsely identified species varied by background: 1 in the soil genome composite, 8 in the gut genome composite, and 12 in the soil isolate composite (Figure 6 B & C). However, these false species accounted for at most 2.59% of the total signal intensity, comparable to the 1.94% of the total intensity attributed to external species by DDA when searching with the gut genome composite. In addition, the KO coverage of the 20 most abundant pathway groups in Mix1 was essentially unchanged, indicating that the functional information captured by these pathways is robust to both missing and spurious organism identifications (Figure 6D). Together, these results demonstrate that this first pass search strategy supports accurate organism identification in complex mixtures and can be combined with downstream functional analysis, thereby enabling DIA-NN’s library-free search on large metaproteomes.

**Figure 6.**
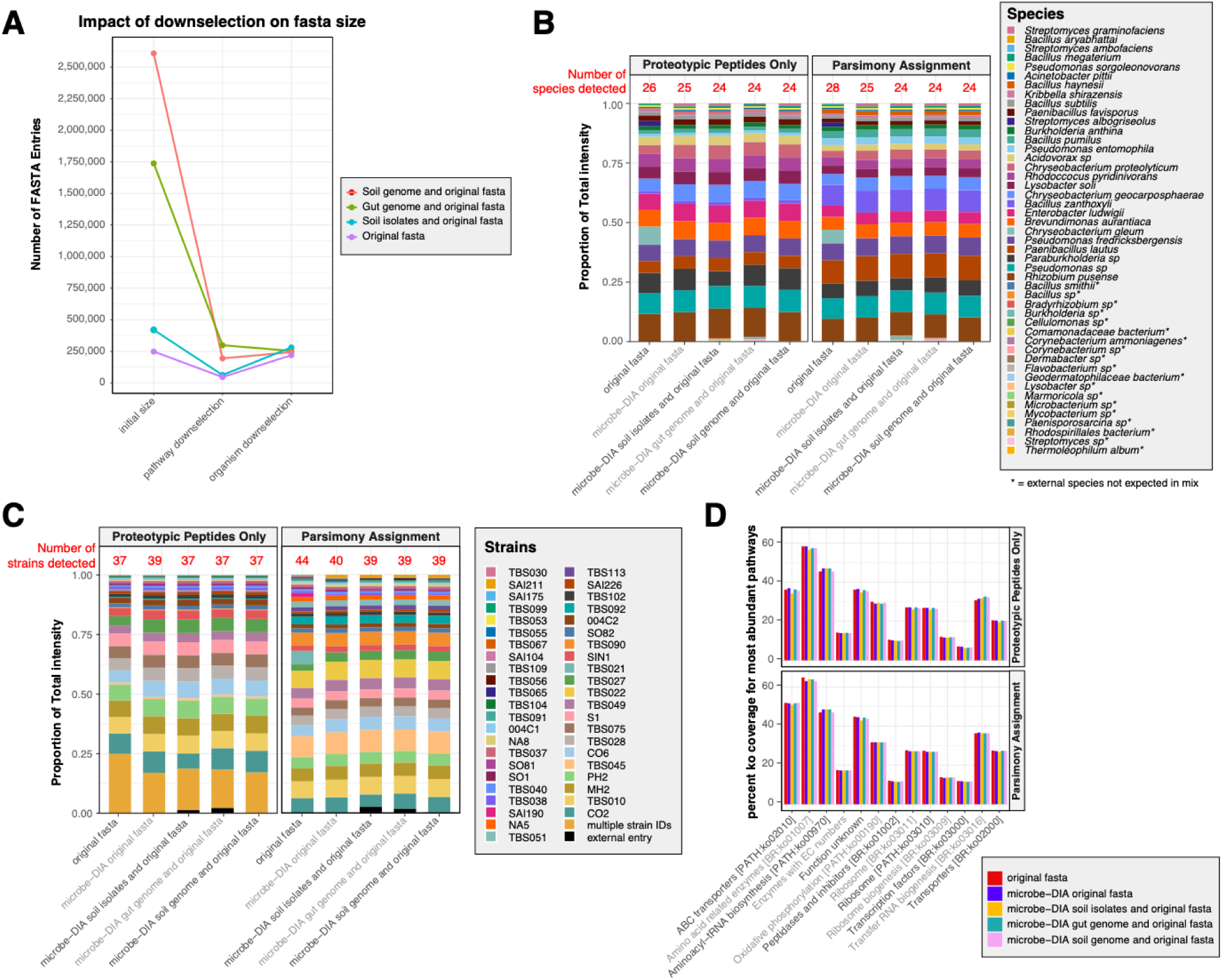
Validation of the two-pass library-free strategy using commonly conserved pathways on the 46-strain DIA Exploris dataset. **(A)** Commonly conserved pathways effectively downselected the test genomes to a manageable search space. After the first and second passes of DIA-NN library-free, the 46-strain dataset’s Mix1 was accurately reconstituted at both the species **(B)** and strain levels **(C),** with a maximum species-level false discovery rate of 2.59% of the total abundance. **(D)** The limited false discovery and missed species do not substantially alter the KO coverage for the most abundant pathway groups, indicating that the functional readout is preserved.

## Discussion

Prior work has shown that data-independent acquisition (DIA) offers clear advantages over data-dependent acquisition (DDA), including improved quantification and reduced missingness, even in the complex case of metaproteomics. However, most studies stop short of defining a complete, metaproteomics-specific DIA workflow that spans from LC-MS/MS data acquisition through to data analysis. A frequently cited obstacle to a fully DDA-independent pipeline is the computational burden of library-free approaches when applied to very large metaproteomic databases, even though these methods have proven effective in more limited, well-defined systems.

Within this context, we systematically evaluated DIA acquisition parameters and found that MS2 window size is a major determinant of DIA identification performance, whereas refinement of the MS1 m/z range has a relatively minor impact on the number of proteins and peptides identified. A smaller MS2 window size appears to be preferable for complex metaproteomic samples, likely because it reduces peptide interferences in the already complex DIA spectra. As DIA scoring algorithms continue to improve, larger MS2 window sizes may enable higher throughput without sacrificing proteome depth. As LC-MS/MS instrumentation evolves, we anticipate incremental increase in throughput in metaproteome analyses, as demonstrated by the substantially higher identification yield on the newer Astral instruments compared to the older Exploris. The comparable number of correctly quantified proteins on the faster Astral instrument suggests that aggressive DIA duty cycles can be adopted without compromising quantitation accuracy. Collectively, these refinements in LC-MS/MS DIA can enable study designs with many more biological replicates, time points or environmental conditions. In the long term, this increase in throughput creates a powerful synergy for the age of AI; by generating massive datasets required for machine learning, our workflow enables the discovery of subtle, complex patterns in microbiome function that were previously beyond our analytical reach.

DIA’s ability to recover information is comparable to deeply fractionated DDA data, but in an order-of-magnitude less instrument time, makes it the practical choice for experiments focused on broad functional profiling and comparative analyses across many samples. However, fractionated DDA data may remain the preferred method when fewer samples are being processed or when cleaner MS/MS for maximal sequence coverage is required, for instance, in novel protein or post-translational modification (PTM) discovery. Future work should focus on characterizing the physicochemical properties of the proteins or peptides uniquely identified by DDA to determine if they represent a specific biological class that is systematically under sampled by DIA. Aside from such targeted objectives, the marginal gain in identifications from extensively fractionated DDA is small relative to the cost in instrument time and sample consumption. Thus, for the majority of modern metaproteomic studies, focused on systems-level comparison, DIA represents the most judicious investment of analytical resources.

A major bottleneck in applying DIA-NN’s library-free mode to real microbiomes is the computational cost of searching against large metaproteomic FASTAs. Without a focused spectral library, the algorithm must consider every possible peptide from the entire database, causing the search space to scale exponentially with the database size. Our MADL two-pass approach partly circumvents this limitation by effectively shrinking the search space while preserving organism-level and functional readouts. The low false organism attribution (<=2.59%) across various sources of background noise, including soil isolates, a human gut metaproteome, and a soil metaproteome, suggests that our approach is robust and scalable.

Despite this success, we observed that the filtering step led to the omission of a few organisms. Specifically, *Streptomyces ambofaciens*, *Streptomyces albogriseolus*, and *Chryseobacterium gleum* were missed in the initial down selected FASTA, while *Bacillus aryabhattai* was missed across all three noise-added conditions. The exclusion of the *Streptomyces* and *Bacillus aryabhattai* may be explained by their high phylogenetic relatedness to other organisms in the community (Supplementary Figure 1), making them difficult to discern from background noise or from other organisms present. This ambiguity was likely amplified by the presence of an unclassified species of *Streptomyces* in the gut metaproteome database. In contrast, the reason for the exclusion of *Chryseobacterium gleum*, which has a medium relatedness with the rest of the organism pool, remains unclear and requires further investigations.

With the computational hurdle of database size effectively removed by our approach, the quality and completeness of the input metagenome now become the primary determinant of analytical success. As such, the use of long-read sequencing to produce high-quality, contiguous metaproteome-assembled genomes (MAGs) are likely to improve the true and false-positive organism attribution by reducing genomic fragmentation and peptide attribution ambiguity. Equally critical are improvements in functional annotation, especially for less-studied organisms, as one of the main goals of metaproteome analyses is to determine the functional activity of bacterial communities. Investments in sequencing and functional sequence attributions will therefore be directly beneficial to both this pipeline and to metaproteomics analyses in general.

Our comparison between proteotypic-only and parsimony-based approaches demonstrates that peptide assignment strategies can substantially increase the number of correctly quantified proteins, especially in DIA. However, the additional complexity this introduces deserves deeper scrutiny, particularly for closely related organisms. Establishing standardized benchmarks to evaluate these peptide-to-protein attribution strategies for organisms with varying relatedness, as we’ve initiated here at the species and strain level, is crucial for ensuring that metaproteomics results are consistent and comparable across studies.

## Conclusions

In conclusion, we establish the <u>M</u>etaproteomic <u>A</u>nalysis with <u>D</u>IA Library-free (MADL) pipeline as a practical and scalable approach for high-throughput metaproteomic analyses. By combining DIA acquisition with a two-pass library-free search, MADL empowers researchers to replace extensive DDA library generation in most studies, dramatically increasing the throughput while maintaining high organismal and functional resolution. While DDA-based strategies will remain essential for specialized applications like de novo protein discovery or the establishment of the role of post-translational modifications in communities, MADL is poised to become a central method for large-scale, quantitative metaproteomics studies that are critical for advancing our understanding of complex microbial systems.

## Supporting information

All suplementary information mentioned in the draft are provided in the Supplemental File.

## Availability of data and materials

The datasets supporting the conclusions of this article are available in the MassIVE repository under accession MSV000102084 at https://massive.ucsd.edu/ProteoSAFe/dataset.jsp?task=0943b6ba567a40e1841baf0c1e3071f8. The R scripts and small relevant files used to generate this analysis are available via archived version TBD-Zendodo of home page TBD. Reuseable code for the Metaproteomic Analysis with DIA-NN Libraryfree (MADL) pipeline is available via archived version TBD-Zendodo of home page TBD.

## Acknowledgements

Thank you to the Pacific Northwest National Laboratory’s Persistent Control Research team and Soil SFA microbial communities’ team, especially to Dr. Rob Egbert, Dr. Ryan McClure, and Andrew Wilson for providing the cultured microbes. Thank you to the Joint Genome Institute (JGI) for generating the soil isolates and soil metaproteome FASTAs we used as external search databases. Thank you also to the Human Microbiome Project for generating the gut metaproteome FASTA we used as an external search database. This research was performed on a project award (10.46936/intm.proj.2023.60905/60008964) from the Environmental Molecular Sciences Laboratory, a DOE Office of Science User Facility sponsored by the Biological and Environmental Research program under Contract No. DE-AC05-76RL01830. Portions of this work were supported by the National Microbiome Data Collaborative with funding from the Genomic Science Program in the U.S. Department of Energy, Office of Science, Office of Biological and Environmental Research (BER) under contract numbers DE-AC02-05CH11231 (LBNL) and DE-AC05-76RL01830 (PNNL). PNNL is operated by Battelle for the DOE under Contract DE-AC05-76RL01830.

