## Supplementary material for "Data Independent Acquisition Pipeline for Microbiome Samples (Microbe-DIA)": All suplementary information mentioned in the draft are provided in the Supplemental File.

Supplementary Information

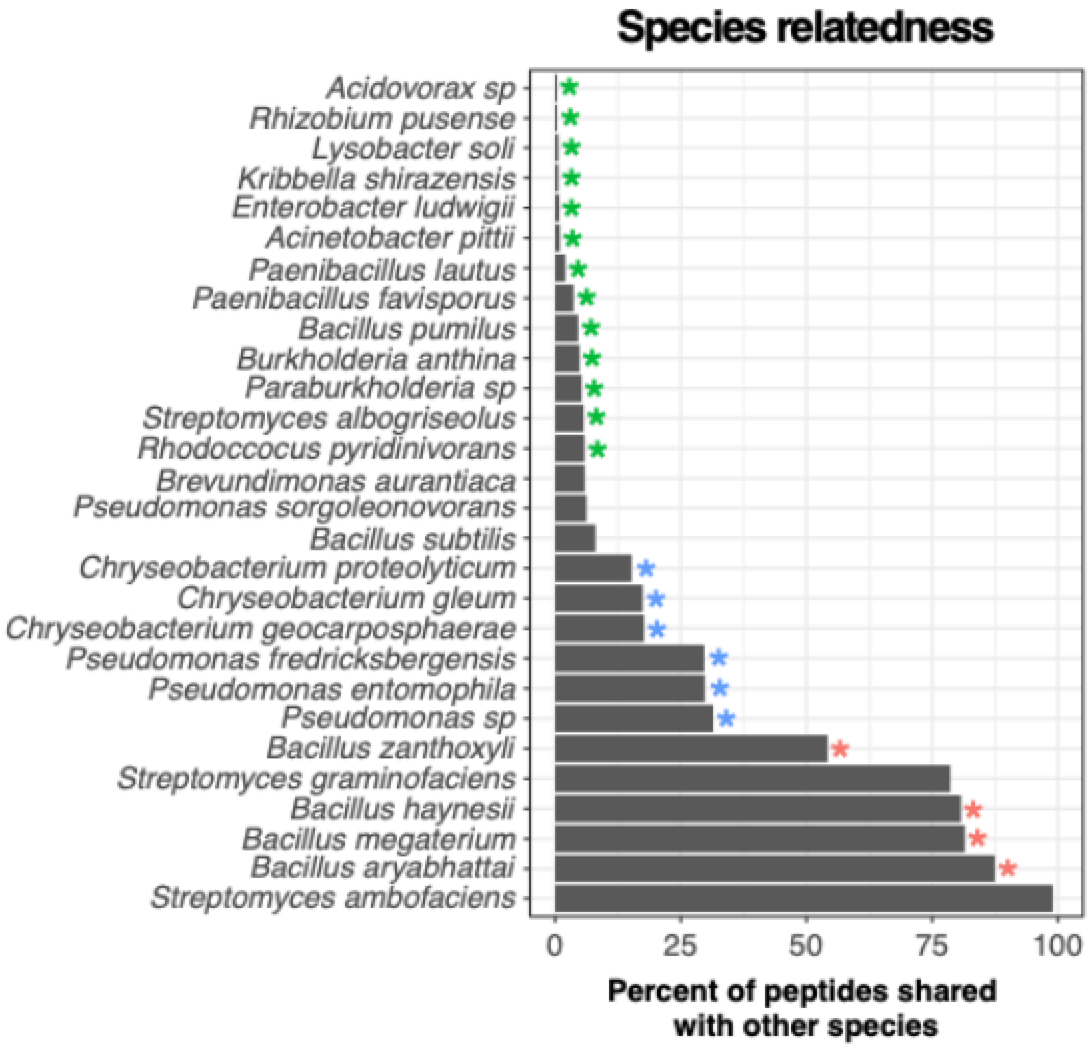


**Supplementary Figure 1**. 24 and 46-strain mixtures were designed using species of varied relatedness to test DIA's performance in progressively complex systems. Relatedness was calculated as the percent of in silico predicted tryptic peptide sequence fragments (of length greater than 10) overlapping amongst available species. Starred species were represented in the 24-strain dataset. Color indicates relative degree of relatedness (green –

low (0.4-5.3%), blue – medium (12-38%), red – high (>50%). All species listed here were represented in the 46-strain mixtures.

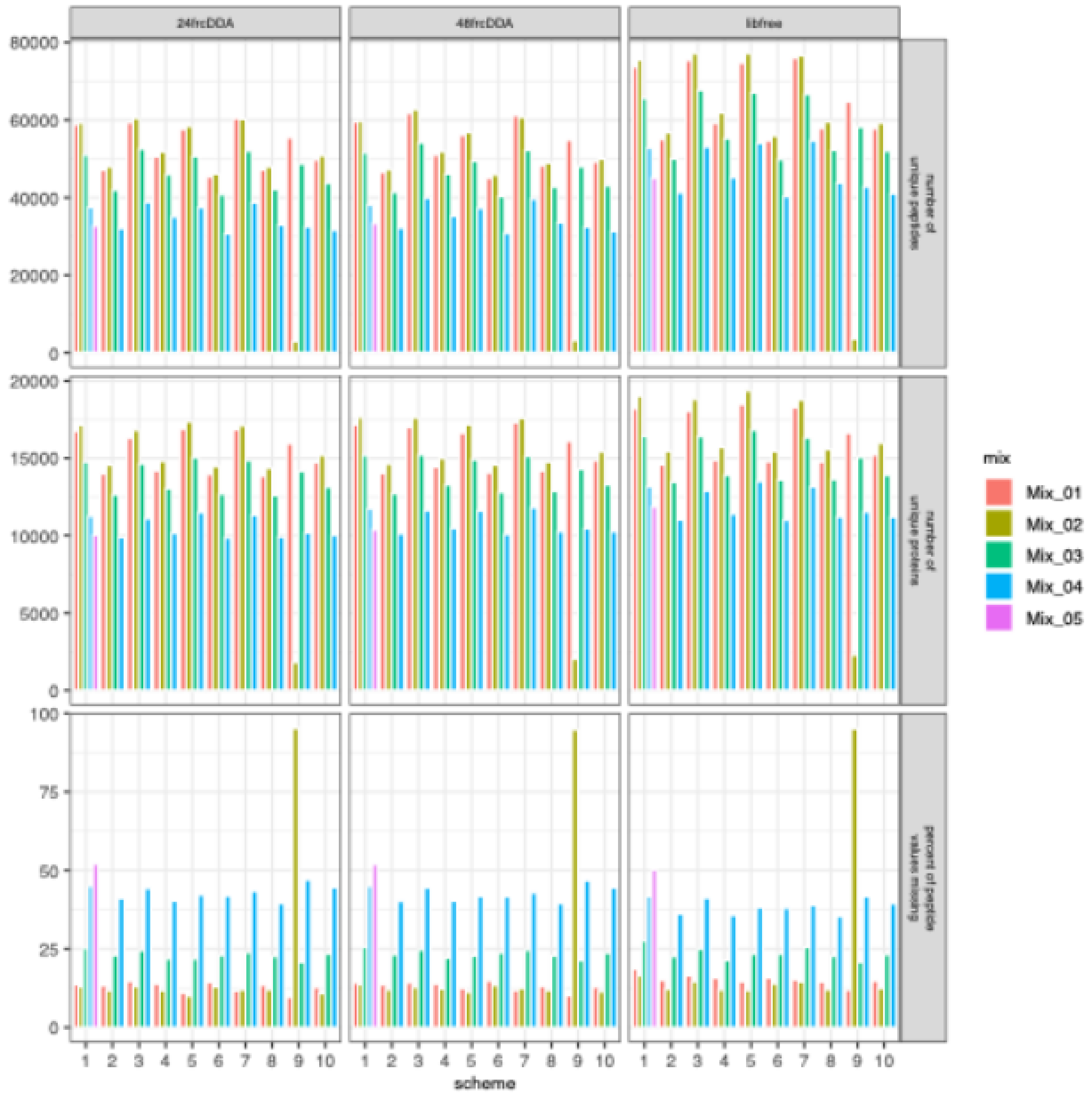


**Supplementary Figure 2**. Comparison of DIA window schemes across all 5 mixtures.

Considering the evident consistency across mixtures, only Mix1 is displayed in Figure 1 so additional elements of each scheme could be displayed, such as MS window size. Note that Mix2 processed in Scheme 9 appears to be an outlier and Mix5 was only analyzed using Scheme 1.

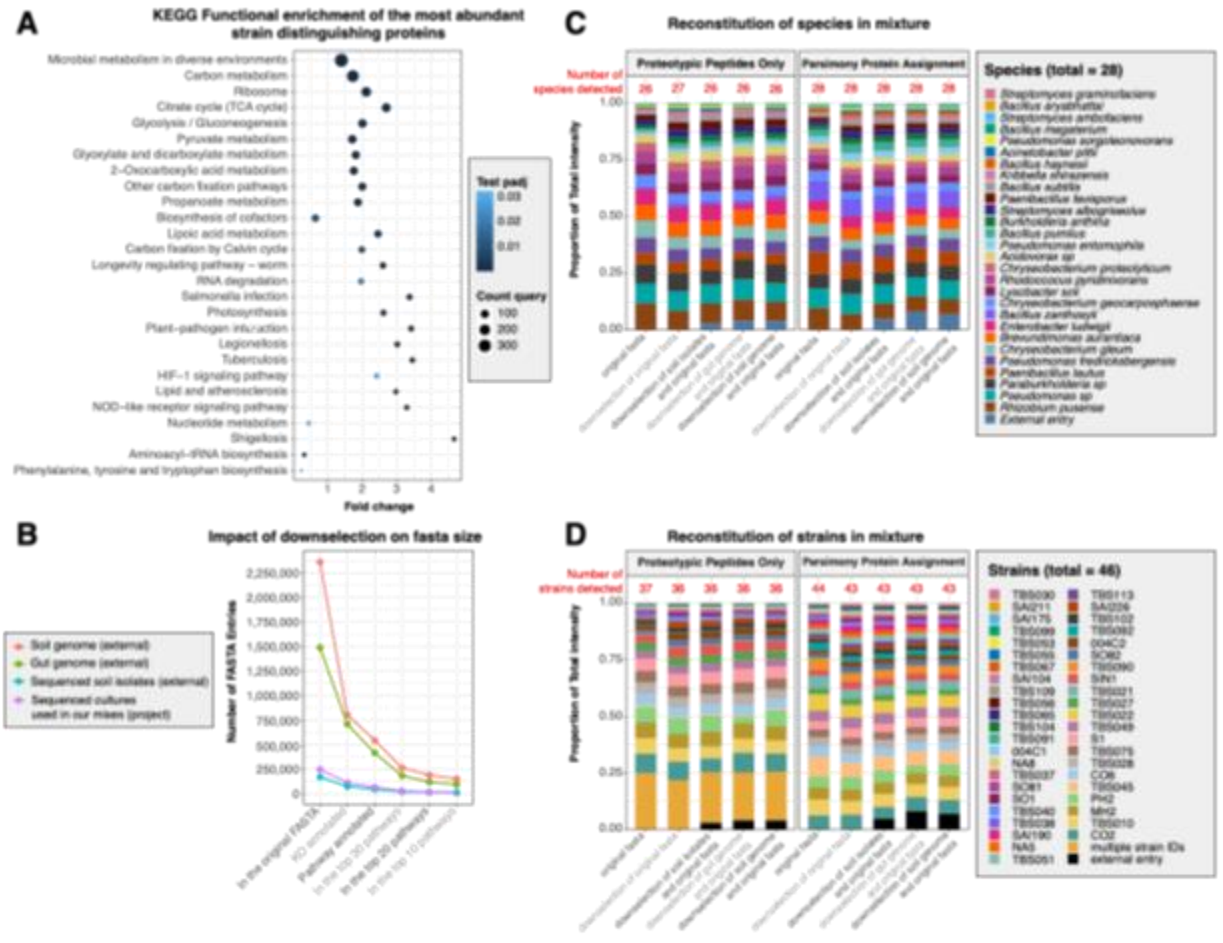


**Supplementary Figure 3**. Initial testing of a two-pass approach to DIA-NN library free. The 46-strain DIA Exploris dataset was used to test whether a two-pass data processing approach can enable DIA-NN's library-free mode on large metaproteomes. **(A)** Pathway enrichment analysis was performed to identify pathways shared amongst the strains, and **(B)** the most significantly enriched pathways were used to down select the metaproteomes to a computationally manageable size. The data were reanalyzed with DIA-NN library-free mode, and the accurately reconstituted Mix1 at both the species **(C)** and strain levels **(D)** demonstrate that signals from the target organisms are retained when searching with the down-selected metaproteome.

**Supplementary Table 1**. Information regarding the FASTA used to search the 24-strain dataset's mixtures, including the dilutions used in sample prep. Italic formatting reflective of taxonomic assignment.

**Supplementary Table 2**. Information regarding the FASTA used to search the 46-strain dataset, including the dilutions used in sample prep. Italic formatting reflective of taxonomic assignment.

**Supplementary Table 3**. Information regarding the expected log2 fold change for the 46-strain dataset. Italic formatting reflective of taxonomic assignment.
